# Fluctuation dominated ligand binding in molten globule protein

**DOI:** 10.1101/2023.04.28.538683

**Authors:** Abhik Ghosh Moulick, J. Chakrabarti

## Abstract

A molten globule (MG) state is an intermediate state of protein observed during the unfolding of the native structure. In MG states, milk protein *α*-Lactalbumin (aLA) binds to oleic acid (OLA). This MG-aLA-OLA complex, popularly known as XAMLET performs cytotoxic activities against cancer cell lines. However, the microscopic understanding of ligand recognition ability in MG state of protein is not yet explored. Motivated by this, we explore binding of bovine aLA with OLA (BAMLET) using all atom molecular dynamics(MD) simulations. We find the binding mode between MG-aLA and OLA using the conformational thermodynamics method. We also estimate the binding free energy using the umbrella sampling (US) method for both MG state and neutral state. We find that the binding free energy obtained from US is comparable with earlier experimental results. We characterize the dihedral fluctuations as the ligand is liberated from the active site of the protein using steered molecular dynamics. The long-live fluctuations occur near the ligand binding site, which eventually transfers towards Ca^2+^ binding site as the ligand is taken away from the protein.

**TOC Graphic:** 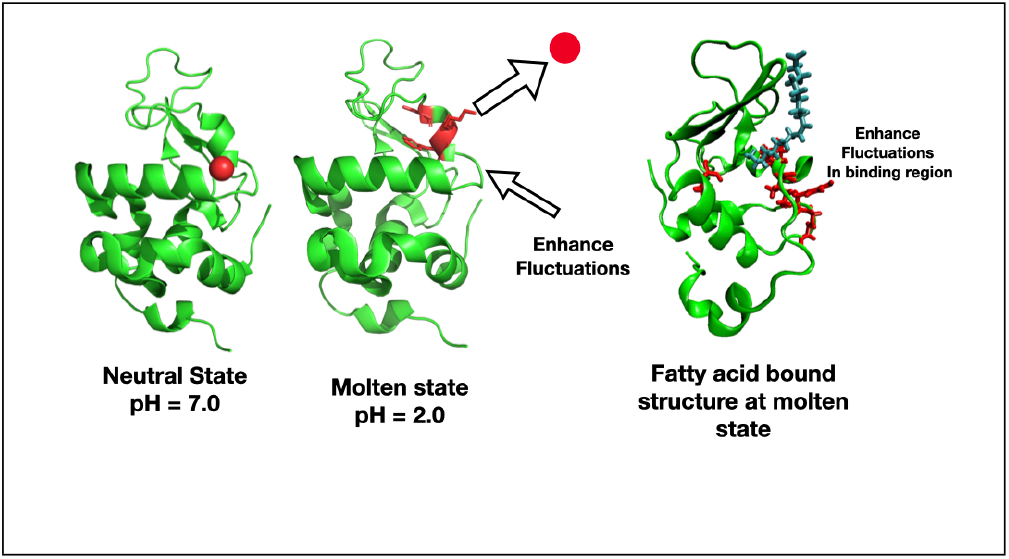

## Introduction

Proteins in a near denaturing condition, also known as molten globule (MG) state, ^1–5^ undergo strong conformation fluctuations^6–9^ while retaining overall secondary structure. In a near denaturing condition, like in a slightly acidic medium, milk protein *α*-lactalbumin (aLA) acts as a carrier of cytotoxic oleic acid (OLA).^10^ This complex is popularly known as XAMLET, namely, X (species) *α*-lactalbumin Made lethal to tumor cells. The participation in cytotoxic activities make XAMLET quite important. The experimental probes, like X-ray crystallography and Nuclear Magnetic Resonance, used to probe structured proteins, can not detect the fluctuating parts in a molten globule protein reliably. Hence, no experimentally probed structure of the XAMLET complex has been reported so far. It may be noted that aLA in its native state does not act carrier of OLA.^11^ Thus, the drug carrier activity of XAMLET is fluctuation driven, whose microscopic mechanism is far from understood.

The protein aLA has a Ca^2+^ binding site, and the crystal structure of Ca^2+^ bound aLA (Holo-aLA) has been reported.^12^ Fig.1 shows the crystal structure of Holo-aLA with crystal water participating in the coordination of Ca^2+^ ion. The protein has *α*-helical (A1-A4) and beta-sheet (B1-B3) domains separated by a cleft (Fig.1). Calorimetric measurements^13^ show that aLA binds with Ca^2+^ ion at the metal binding loop connecting two domains. The binding site of Ca^2+^ is formed by the carbonyl oxygen of Lysine (LYS)79 and Aspartate (ASP)84, the side chain carboxylates of ASP82,ASP87, and ASP88. In addition, the crystal water molecules take part in coordination. At acidic pH, protons compete with Ca^2+^ ion for the carboxylate oxygens and disturb the coordination of Ca^2+^ ion. Hence, Ca^2+^ gets depleted in the MG-aLA state. In an earlier study ^14^ we show that protein at MG state behaves as external condition-induced disordered protein which undergoes conformation fluctuations similar to those in intrinsically disordered protein (IDP).

It is observed that bovine Holo-aLA is unable to bind OLA. But, MG-aLA binds to OLA with binding free energy −9.45 kcal/mol. ^11^ Experimental studies ^15^ suggest that the putative binding site of OLA lie between the A1 and A2 helices and the inter-facial cleft (See, Fig.1).

**Figure 1:**
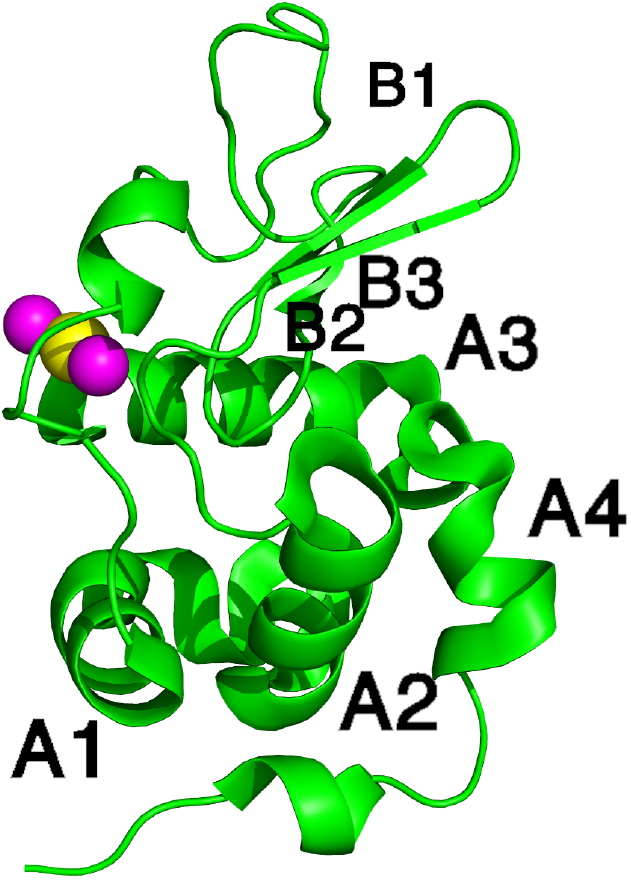
Initial crystal structure of Holo *α*-lactalbumin protein. Ca^2+^ ion is shown in yellow, and crystal water participating in the coordination of the ion is shown in magenta. The secondary structure element of the alpha-helical (A1–A4) and the beta-sheet (B1–B3) domains are marked.

It has been proposed that OLA stabilizes the MG state in XAMLET. ^16^ However, nothing is known microscopically about ligand recognition by proteins in the MG state. Motivated by this, we explore binding of bovine MG-aLA with OLA using the all atom molecular dynamics(MD) simulations. Our objective is to understand how the microscopic fluctuations inherent in MG state dominate the binding residues of the ligand.

We consider in particular the bovine aLA, namely, BAMLET complex. Since there is no experimental data available on the binding mode, we build up a model BAMLET with the help of thermodynamic cost of conformation changes, calculated from the dihedral fluctuations of a protein over constant pH molecular dynamics (CpHMD) simulation^17^ trajectory in an acidic medium with respect to that in neutral medium.^18–20^ We use the model BAMLET to further molecular dynamics (MD) simulations along with umbrella sampling to understand the stability of BAMLET within our model. We observe a stable BAMLET complex with binding energy is −8.3 kcal/mol, comparable with earlier experimental results. ^11^ This also validates our model BAMLET structure. We perform steered molecular dynamics to generate conformations for various distances of OLA from protein to understand how the fluctuations govern kinetics of OLA binding in the context of our model BAMLET. We identify the fluctuations in the system using the dihedral-based PCA, dPCA+, and machine learning (ML) techniques.^21–24^ We observe that the essential coordinates(EC) in the conformation fluctuations^14^ belong to some of the ligand-binding active residues. These degrees of freedom undergo a small change in conformation free energy in BAMLET as compared to MG-aLA conformations. This suggests that the OLA binding is strongly dominated by fluctuations. As the ligand goes further away, the EC has shifted towards the residues of Ca^2+^ binding region, suggesting that the Ca^2+^ binding residues have allosteric control on the ligand binding in agreement with the earlier works. ^14,19^

## Methods and analysis

### Constant pH molecular dynamics

The protein considered in this study is Bovine-*α*-lactalbumin in both Holo (with Ca^2+^ ion, RCSB PDB ID: 1F6R) and apo (without Ca^2+^ ion, RCSB PDB ID: 1F6S) ^12^ form. Both crystal structures have 6 identical chains, out of which we choose only chain A in the analysis. The initial structure is shown in Fig.1. To maintain the pH of the system, we use the protocol of CpHMD simulation in explicit solvent suggested in Ref. ^25^ and used in our earlier work^14^ keeping the number of particles (N), volume (V) and temperature (T) fixed. We perform constant pH at pH=2 using AMBER^26^ simulation package. We use aspartate (ASP) and glutamate (GLU) for titration at pH=2.0. Histidine (HIS) is excluded due to the high acidic pH (=2). HIS has pKa value 6.5 hence Under neutral conditions, pH = 7, only HIS is considered for titration. The details of constant pH simulation of apo-aLA at pH=2.0 (MG-aLA) and normal MD of Holo-aLA is discussed in Ref.^14^

### Conformational thermodynamics

The free energy and entropy associated with conformational changes are computed from the histograms of the dihedral angles, as reported earlier.^18^ The conformational free energy and entropy cost for an individual protein residue can be estimated by taking the sum of the contribution of free energy and entropy of each dihedral in that residue. We compute this with our in house tools, which can be obtained from the GitHub (**https://github.com/snbsoftmatter/confthermo**). Residues having Δ*G,T*Δ*S >* 0.0 are identified as destabilized and disordered residues. These residues are active in ligand binding. This yields the binding mode between the ligand and the protein.

### Protein-ligand complex preparation

#### Clustering and docking

As no crystal structure is available for MG-aLA, we use K-means clustering ^27,28^ to identify representative structure over the trajectory of CpHMD run at pH=2.0. In molecular simulation, clustering represents the grouping of similar conformations together. Similarity is determined by a distance metric, where the smaller distance represents more similar structures. Here, coordinate root-mean-square deviation (RMSD) is used as distant metric. Clustering is computed using CPPTRAJ^28^ tools of AMBER.^28^ Next, we dock protein structure obtained from clustering of MG-aLA using the HADDOCK. ^29,30^ We dock the ligand using conformationally destabilized and disordered residues of cleft region as bias. The docking protocol consists of following stages, at first a rigid body energy minimization and then MD base refinement process provides model structures. Basis on Z score, therefore, the top cluster is chosen. Details of docking are in SI Table.1. The OLA molecule is taken from the crystal structure of liver fatty acid binding protein–oleate complex (PDB ID: 1LFO). We use similar protocol to construct the model for Holo-aLA-OLA complex.

#### MD simulations of protein-ligand complex

The MD simulations of the docked complex are performed using GROMACS^31^ simulation package with Amber99Sb force field^32^ and TIP3P^33^ water model. The GROMACS parameter for OLA are generated using Antechamber and Acpype. Antechamber parametrizes the molecule using general AMBER force field (GAFF).^34^ Acpype is python interface to antechamber, which provide GROMACS topologies.^35^

AMBER CpHMD method provides fraction of time titrable residues protonated during the simulation. SI Table 2 shows fraction of time titrable residues remain protonated during simulations. Based on this, we set protonation state of titrable residues in GROMACS.

We further perform a total of 2 sets of simulations : (i) MG-aLA-OLA and (ii) Holo-aLA-OLA complexes. At first, systems are immersed in a cubic box of dimension 7.221×7.221×7.221 nm^3^ and 7.725×7.725×7.725 nm^3^ respectively. 11 Cl^-^ ion is required for protein-ligand complex at MG state. Holo-aLA required 6 Na^+^ ion to neutralize the complex structure. Minimization is done for 50,000 steps using the steepest descent algorithms. All bonds in the protein are constrained using the LINCS algorithm. Equations of motion are integrated using leap-frog algorithm with an integration time step of 2fs. Systems are equilibrated through 2 steps (NVT and NPT) using position restraints to heavy atoms. The NVT and NPT equilibration is carried out at 300K Temperature and 1 Bar pressure. Afterward, the full all-atom simulations are performed for 1 *µ*s with 2 femtosecond time step integration employing periodic boundary conditions in all directions.

#### Steered MD and Umbrella sampling(US) simulation

The equilibrated structure of MG-aLA-OLA complex is selected for the US study. The initial system is prepared using GROMACS software and protein-OLA complex at MG state is made parallel to the Y axis. The dimension of the box is 6.53×11.95×4.34 nm^3^. The prepared box is solvated, neutralized, and minimized following the previous simulation protocol. The box is further equilibrated at a specific temperature (NVT) and specific pressure (NPT) similar to the previous MD simulation setup. The US method initiates with steered molecular dynamics(SMD) method using the distance between center of mass as reaction coordinates. The OLA molecule is pulled from binding residues of protein towards the bulk solvent for 500 ps with 1000 kJ/mol-nm force. OLA molecule is pulled at the rate of 0.01 nm per ps. Snapshots are saved at each pico second during the course of pulling, hence total of 500 configurations are generated. We extract 22 frames from the SMD trajectory to prepare the umbrella sampling windows, where the distance between each configuration is 0.2 nm. These configurations serve as the initial structure of US simulation and each frame is independently equilibrated, performing NPT equilibration for 100 ps. Next, MD is performed for each individual configuration. The potential of mean force (PMF) is calculated from equilibrated trajectory of the US simulation using the weighted histogram analysis method (WHAM),^36^ included in GROMACS. One can calculate binding energy from PMF curve by subtracting the PMF value at the position of the ligand at the maximum distance from the PMF value at the position of the ligand at the minimum distance from the protein. We also run separate umbrella sampling where complex structure obtained from equilibrium MD trajectory is used as the initial configuration. We calculate the error bar multiple PMF obtained from 5 independent umbrella sampling simulations. In total, we performed 4.3 *µ*s of US simulation. We also perform US simulation for Holo-aLA and OLA following similar protocol.

### Analysis

#### Identification of essential coordinates (EC)

We identify essential coordinates of the system for various positions of OLA molecules from aLA at MG state. For this, we used SMD techniques as discussed earlier, where the distance between the center of mass between protein and ligand is used as the reaction coordinate. The essential coordinate is identified using the method described by Brandt et al.^21^ The method is used earlier to identify essential coordinates in the MG state of aLA (MG-aLA) in Ref.^14^

#### Dynamical cross-correlation analysis

We calculate the cross-correlation function *C*(*i, j*) between Δ*r*_*i*_ and Δ*r*_*j*_, the displacement vectors of i-th and j-th *C*_*α*_ atom of the protein.^37,38^ During this analysis, the trajectory at *t* = 0 is considered as a reference structure, and all other structures are aligned with respect to that reference structure. The Bio3D suite of R programming packages^39^ is used for this calculation.

#### Radial distribution function

We calculate the radial distribution function (rdf), g(r) to understand the arrangement of water around the protein surface. The rdf is calculated using the following equations:

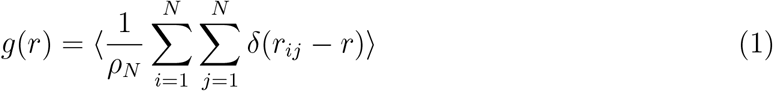

where denotes the total number of water molecules, *ρ*_*N*_ is the particle density and the angular bracket represents average over time origins.

## Results and discussion

### Model BAMLET structure and validation

Since no experimental data are available on the binding mode, we build up a model BAMLET structure. Recent studies show that the free energy and entropy costs associated with conformation changes computed based on the dihedral fluctuations of a protein over simulated trajectories in different conformations. ^18–20^ Conformationally destabilized residues with increasing free energy and disordered residues with increasing entropy are shown to participate in ligand binding.^19^ We simulate the protein at neutral pH and pH=2 using the constant pH molecular dynamics (CpHMD) technique^17^ and compute conformational thermodynamics data in the MG-alA state with respect to the Holo-aLA state. The changes in free energy and entropy consist of contributions due to the change in dihedral distributions of backbone dihedral angle *ϕ, ψ*, and all the side chain dihedral angle *χ*_*i*_, *i* = 1, .., 5. Fig.2(a) shows data for changes in conformation free energy (Δ*G*) and entropy (*T*Δ*S*) of the residues in the cleft region. We observe that all the residues are conformationally destabilized and disordered. The total change in Δ*G* and Δ*S* of the binding region of aLA at MG-aLA conformations over Holo-aLA conformations are 26.20 kJ/mol and 68.84 kJ/mol. This set of residues consists of a large number of hydrophobic residues and a few basic residues, listed in the SI Table 3. We consider residues near the cleft region to be prone to binding, in agreement with the earlier study, ^15^ as active residues to dock OLA to a representative structure of the protein in the MG state as initial model BAMLET structure.

**Figure 2:**
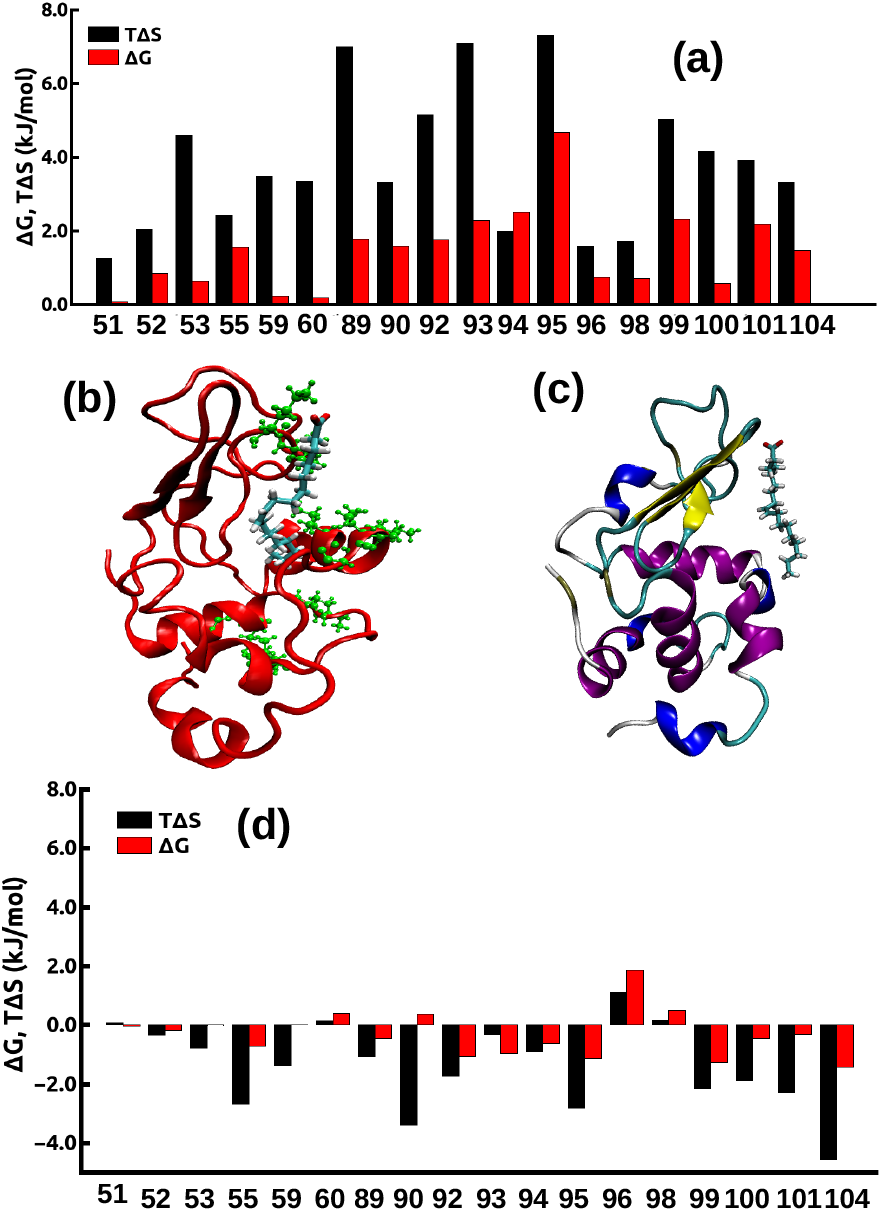
(a) Conformational thermodynamics changes at molten globule state with respect to the neutral state, (b) Equilibrated structure of BAMLET complex. Hydrophobic tail of the OLA goes into the cleft region. Residues of aLA involved in forming binding interface with OLA are marked in green. (c) Equilibrated structure of Holo-aLA-OLA complex. OLA remains outside the protein cleft region all over the simulation, (d)Conformational thermodynamics change of active residues at BAMLET conformations with respect to MGaLA conformations.

We refine the model structure, subjecting the initial structure to MD simulations. An equilibrium structure from MD trajectory of BAMLET complex is shown in Fig.2(b). We identify binding interface in BAMLET complex based on minimum distance between (*d*_*min*_ *<* 4Å) protein *C*_*α*_ atom and main chain carbon atom of OLA. Residues, responsible for forming the binding interface, are marked in green color over the protein in equilibrated structure in Fig.2(b). The long hydrophobic tail of OLA extends towards the interface cleft. We find that interface consists of residues, like LYS58, ILE59, ILE95, LYS98, VAL99, GLY100 of cleft and ARG10, LYS13, LYS16, GLY17, LEU23 of A1-A2 helices. This is in agreement to the experimental studies, suggesting that the putative binding site of OLA lies between the A1 and A2 helices and the interface cleft.^15^ Fig.2(c) shows an equilibrium structure from MD trajectory of Holo-aLA-OLA. It shows that OLA does not bind into the cleft region of Holo-aLA, as observed in experiment. ^11^

We calculate binding energy between protein-ligand using umbrella sampling techniques (Details in Method section) where the reaction coordinate, *ξ* is the center-of-mass (COM) distance between the ligand and the protein. Fig.3(a) shows the PMF curve of the proteinligand complex for BAMLET. The binding energy of the protein-ligand complex is approximately −8.3 kcal/mol, which is comparable with earlier experimental observations. ^11^ In case of Holo-aLA the US method shows much lower binding energy of −2.4 kcal/mol (data not shown here). This also validates our model for BAMLET.

**Figure 3:**
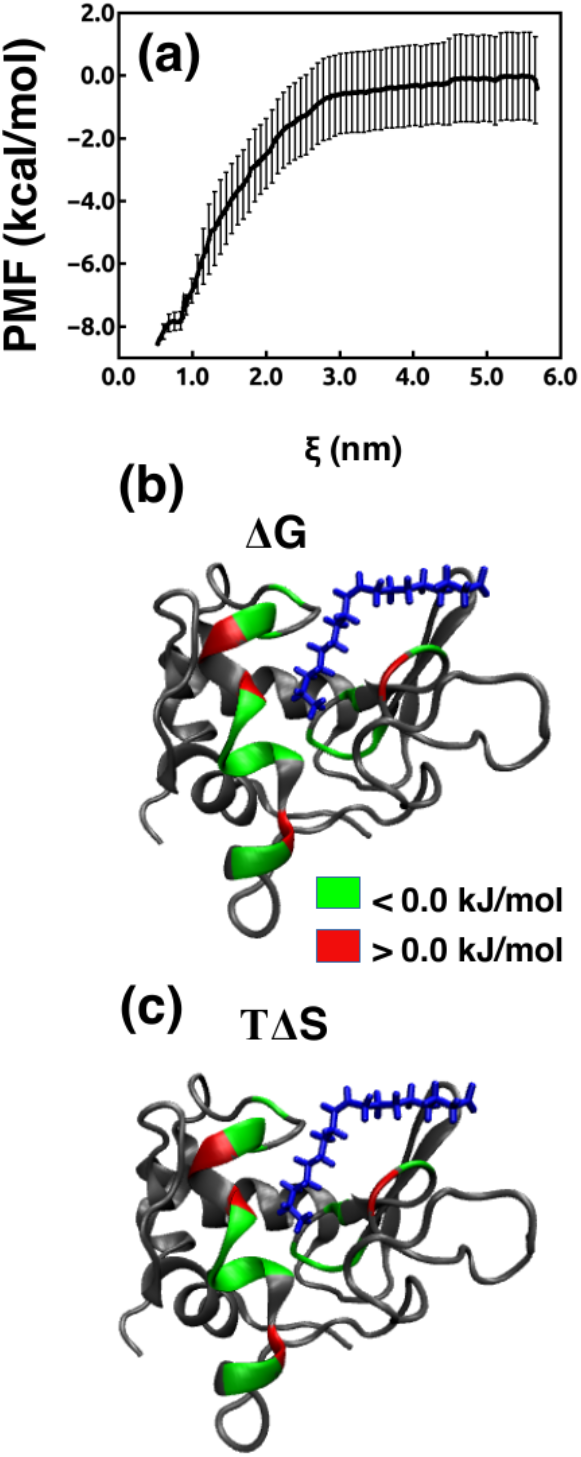
(a) PMF curve of the protein-ligand complex obtained from umbrella sampling method for BAMLET conformations, (b) Δ*G* (c) *T*Δ*S* of active residues in equilibrated structure. The green color shows stabilized/ordered residues, and red corresponds to destabilized/disordered residues.

Next, we check the conformation free energy Δ*G* and entropy *T*Δ*S* costs of the active residues in BAMLET complex with respect to MG-aLA, shown in Fig.2(d). We find that all binding residues have Δ*G <* 0, except TRP60, MET90, LEU96, and ILE98. Those residues are marked in red over the equilibrated structure of BAMLET in Fig.3(b). Most of the active residues are ordered except TRP60, LEU96, and ILE98. Fig.3(c) shows entropy *T*Δ*S* costs of those active residues. The residues marked in green color are entropically ordered, and marked in red color are entropically disordered. All binding residues are *T*Δ*S <* 0 except

TRP60, LEU96 and ILE98 are entropically disordered. The total change in Δ*G* and Δ*S* of the binding region are −5.47 kJ/mol and −24.8 kJ/mol respectively. Thus, the overall binding region of aLA becomes stabilized and ordered due to OLA binding. This gives microscopic justifications to the proposal of OLA-induced stability of MG-aLA. ^16^

### Fluctuations in BAMLET

Now we consider the microscopic fluctuations in BAMLET. We compute to this end the dynamic cross correlation map (DCCM) C(i,j) ^37,38^ between protein residues, (see in the method section) to reveal correlated motions involved in the recognition of ligand. ^38^ Fig.4(a) shows binding region of the protein at MG state in color, where green corresponds to hydrophobic and basic residues of A1-A2 region and red represents hydrophobic and basic residues of cleft region. We compare DCCM map for two different cases: (i) MG-aLA (Fig.4(b)), and (ii) BAMLET complex (Fig.4(c). In MG-aLA, the residues of ligand binding region are negatively correlated(Fig.4(b)). The region of interest is marked within the box. It suggests that putative binding residues near the cleft region (residues within 50 to 100, marked in red in Fig.4(a)) shows anti-correlated behavior with residues belongs to A1-A2 region (residues within 1 to 40, marked in green in Fig.4(a)). This anti-correlated motions between residues of protein decrease in BAMLET (Fig.4(c)). Thus, the long ranged anti-correlated motions between protein segments are involved in ligand binding.

**Figure 4:**
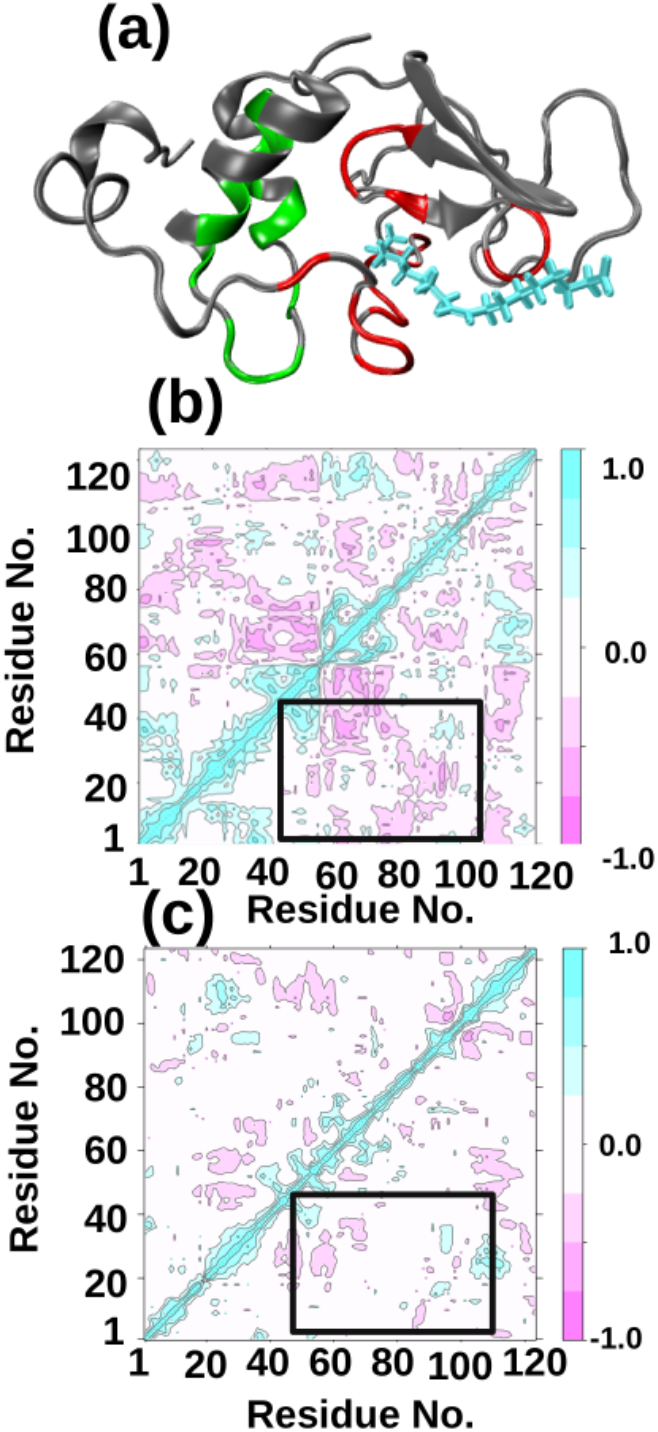
(a) Binding region of the protein at MG state in color, where green color corresponds to hydrophobic and basic residues of A1-A2 region and red color represents hydrophobic and basic residues of cleft region. Dynamic cross correlation map (DCCM) for two different cases, (b) MG-aLA and (c) BAMLET complex.

Next, we characterize the fluctuations in terms of microscopic conformational variables, like the dihedral angles. We construct the free energy landscape by computing population(H_i_) base free energy as a function of the i-th dihedral principal components. Earlier results^14^ show that metastability is enhanced in MG-aLA state as compared to Holo-aLA. Fig.5(a)-(c) shows free energy landscape (FEL) of principal components (PC1 to PC3) in BAMLET. We observe that the meta-stability changes in presence of OLA 5(a)-(c). Along PC1-PC2, the metastability is reduced in BAMLET conformations as compared to MG-aLA conformations reported earlier. ^14^ Along, PC3 metastability completely vanishes in BAMLET, whereas for MG-aLA metastability still present.^14^ Reduction in metastability suggests that at BAMLET conformations, protein has more stable state as compared to MG-aLA conformations, which is consistent with the conformation thermodynamics data of BAMLET.

**Figure 5:**
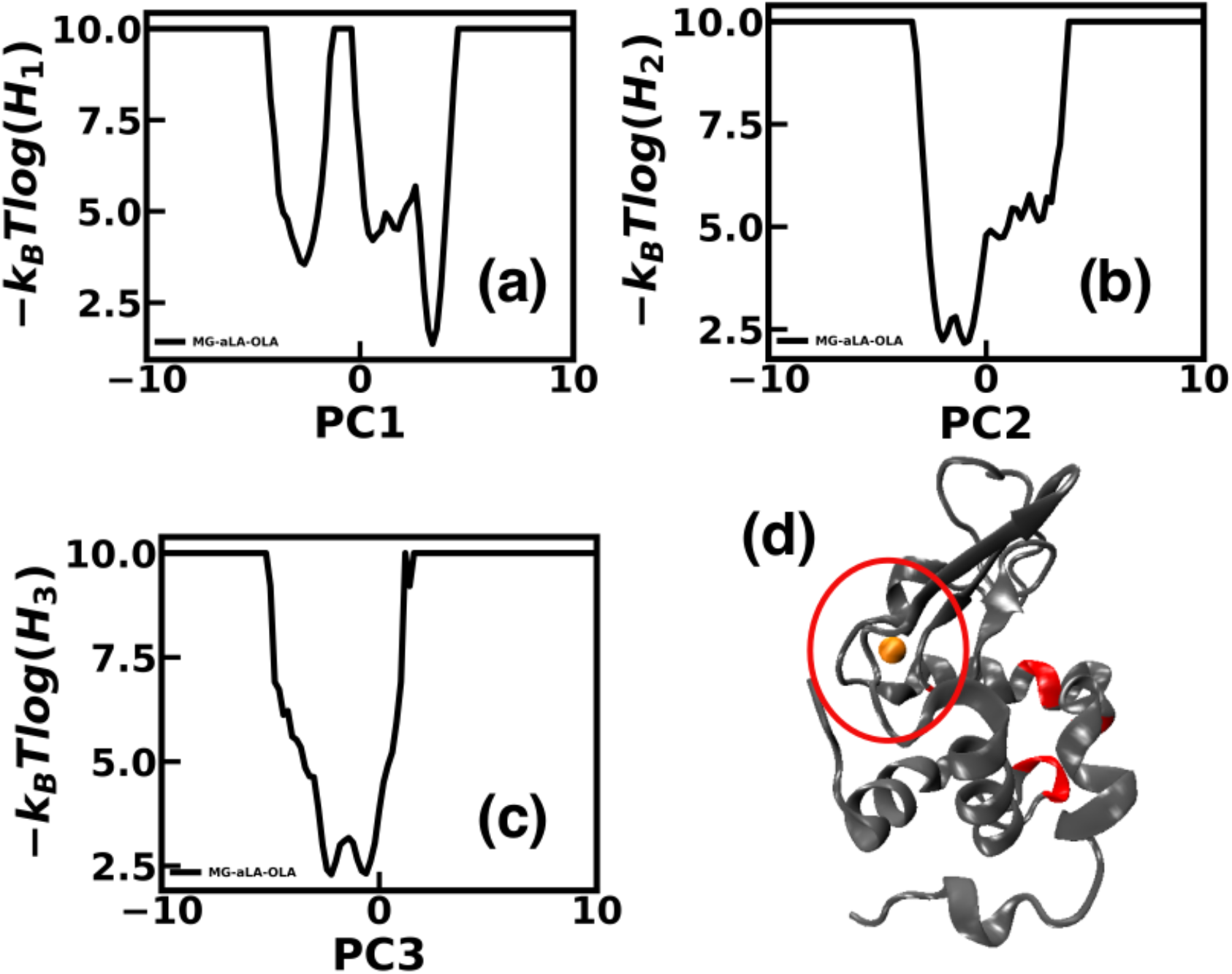
Free energy landscape obtained from dPCA+ along (a) PC1, (b) PC2, (c) PC3 for BAMLET. Y-axis represents negative log of population of PCs (H), (d) Residues having essential coordinate in BAMLET complex.

We identify essential coordinates of the system using clustering and supervised machine learning-based analysis.^21^ Residues having the essential coordinates in protein are colored (red) over the crystal structure in Fig.5(d). The Ca^2+^-binding region is marked within the red circle. ECs are also tabulated in Table.1. The first element of the column denotes the most EC which is obtained in the final iteration of the XGBoost removing all other coordinates. We marked ECs in bold which are either identified as active residue for OLA from conformational thermodynamics analysis. In BAMLET, ECs mostly belong to the active residue for ligand binding such as ILE95 (*ϕ*95), GLY100 (*ϕ*100), LEU52 (*ψ*52), LYS94 (*ψ*94 and *ϕ*94) and ILE101 (*ψ*101). Here, we denote the EC within bracket after each residue by the dihedral angle and residue number. *ϕ*95 acts as the most essential coordinate. This suggests that in the complex, the fluctuations of the system mostly governed by some of the binding residues. We already have discussed in earlier work^14^ that at MG-aLA conformations, EC belongs to *ψ*80 and *ϕ*79 i.e. near Ca^2+^ binding region in stark contrast to the fluctuations in BAMLET. Since the fluctuation scenario is entirely different in BAMLET compared to MG-aLA, we examine the time dependent changes in microscopic fluctuations in response to the approach of the ligand. To this end, we carry out all-atom MD simulations on various configurations with fixed *ξ* obtained during the steered molecular dynamics simulations. We show the FEL along PC1 for different configurations for different *ξ* in Fig.6(a). The figure shows metastability for different *ξ*. Fig.6(b)-(d) shows the residue having essential coordinates in protein for different *ξ* using similar color code in Fig.5(d). Table.1 shows that essential coordinates and corresponding amino acid name for different *ξ*. We mark ECs by star in each column which belongs near the Ca^2+^ binding region.

**Table 1:** Essential coordinate for 4 different cases. The first entry of each column represents the most essential coordinate. EC participates in ligand binding and interface formation between protein and OLA are marked as bold in each column. Residues near the Ca^2+^ binding region are marked with star.

| Complex | | $\xi = 1.34 \text{ nm}$ | | $\xi = 1.93 \text{ nm}$ | | $\xi = 2.5 \text{ nm}$ | |
| --- | --- | --- | --- | --- | --- | --- | --- |
| EC | Residue Name | EC | Residue Name | EC | Residue Name | EC | Residue Name |
| ϕ**95** | **ILE** | ϕ63 | ASP | ϕ39 | GLN | ψ**17** | **GLY** |
| ψ18 | TYR | ϕ**23** | **LEU** | *ϕ84 | ASP | ψ**13** | **LYS** |
| ϕ**100** | **GLY** | ψ107 | HIS | ψ54 | GLN | ψ24 | PRO |
| ϕ20 | GLY | ψ**13** | **LYS** | ψ64 | ASP | *ψ81 | LEU |
| ϕ19 | GLY | *ϕ88 | ASP | ψ91 | CYS | ψ40 | ALA |
| ψ**52** | **LEU** | ϕ49 | GLU | ψ39 | GLN | ϕ24 | PRO |
| ψ**94** | **LYS** | ϕ54 | GLN | ϕ56 | ASN | ϕ**13** | **LYS** |
| ψ**101** | **ILE** | ϕ24 | PRO | ϕ64 | ASP | *ϕ87 | ASP |
| ψ19 | GLY | ϕ56 | ASN | ψ**53** | **PHE** | ϕ25 | GLU |
| ϕ**94** | **LYS** | ϕ**53** | **PHE** | ψ32 | HIS | ψ**55** | **ILE** |

For *ξ* = 1.34 nm, the most EC is ASP63 (*ϕ*63) near one of the binding residues, TRP60. The other ECS, like LUE23 (*ϕ*23),LYS13(*ψ*13) and PHE53 (*ϕ*53) also belong to the region near the active site for OLA binding. On the other hand, ASP88 (*ϕ*88) belongs to the Ca^2+^ binding region (Fig.6(b)). For *ξ* =1.93 nm (Fig.6(c)), the most EC belongs to GLN39 (*ϕ*39). We find residue like PHE53 (*ψ*53) as one of the EC which acts as active residue for OLA binding. We also find some other residues have EC near the vicinity of putative binding site for OLA identified using conformational thermodynamics, like: GLN54 (*ψ*54), CYS91 (*ψ*91), ASN56 (*ϕ*56). Both dihedral *ϕ, ψ* of residue ASP64 near binding residue TRP60, act as EC. On the other hand, ASP84 (*ϕ*84) acts as one of the EC which belongs near the Ca^2+^ binding loop. For *ξ* =2.5 nm, the most EC is shifted to GLY17(*ψ*17) shown in Fig.6(d). Apart from some active site residues holding the ECs, we find that ASP82 in the vicinity of LEU81(*ψ*81)and ASP87(*ψ*87), directly participating in Ca^2+^ coordination, also bear ECs. Thus, there is gradual shift of the ECs from ligand binding residues when ligand is in complex to Ca^2+^ binding loop region as ligand-protein distance increased. This suggests allostery between OLA binding and the Ca^2+^ binding loop residues.

**Figure 6:**
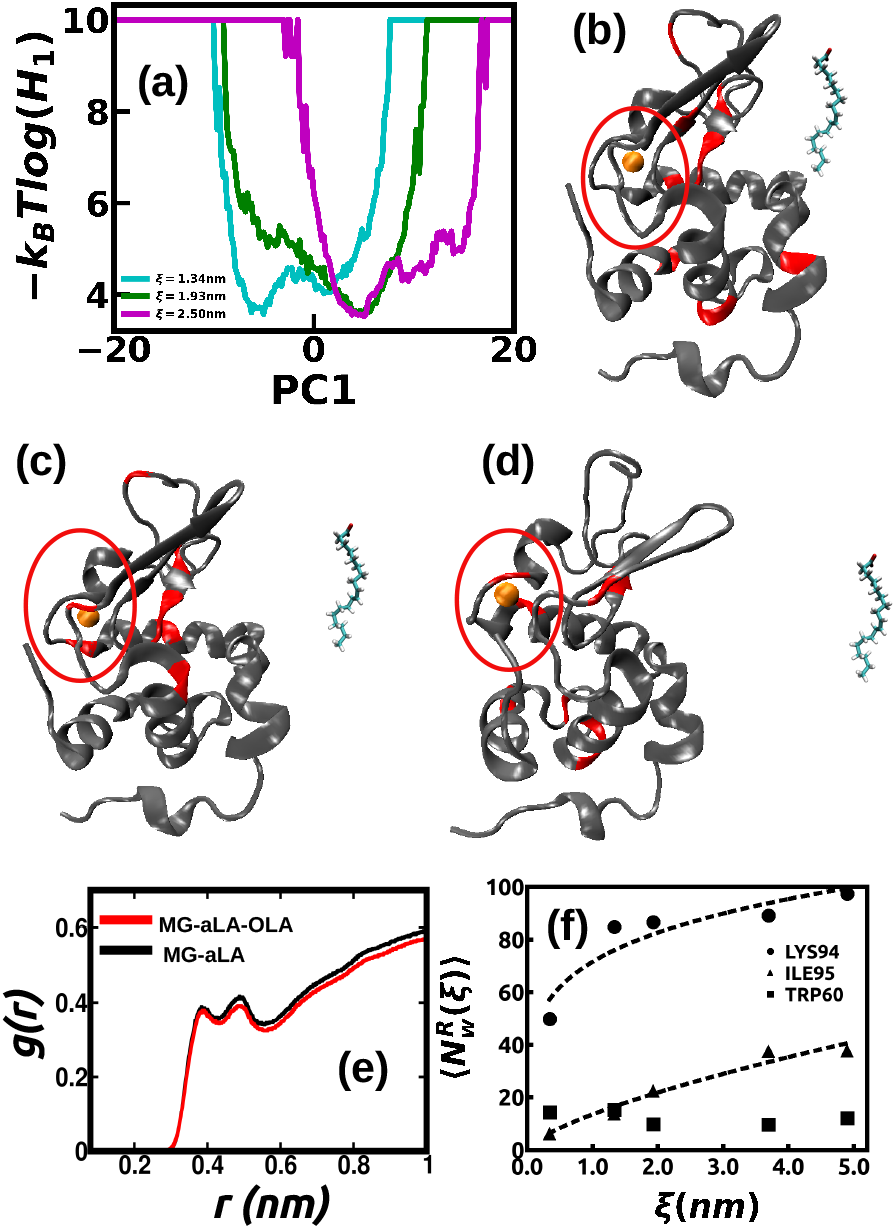
(a) Free energy landscape along PC1 obtained from dPCA+ for different *ξ*. Essential coordinates are colored in red over crystal structure of aLA for (b) *ξ* = 1.34 nm, (c) *ξ* = 1.93 nm, (d) *ξ* = 2.5 nm. The ligand is shown in the figure. We also show Ca^2+^ binding region of protein aLA along with Ca^2+^. (e) Radial distribution function, g(r) of water molecules as a function of distance from the protein at MG-aLA and BAMLET conformations (f) Average number of water molecule 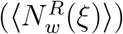 around ILE95 (triangle), LYS94 (circle) and TRP60 (rectangle). Fitted lines are shown for ILE95 and LYS94.

Since hydrophobic regions participate in binding in BAMLET, we examine the behavior of water around the binding residues. We calculate the radial distribution function (g(r)) of water molecule around *C*_*α*_ atom of amino acid residues, where r is defined as the distance between *C*_*α*_ atom of an amino acid and oxygen atom of a water molecule. Fig.6(e) shows g(r) plot for MG-aLA (black) and BAMLET (red). The peak is mostly unaffected due to complex formation in BAMLET as compared to MG-aLA. We calculate average water number 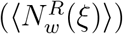 around the R th residue for different *ξ* within the first peak of g(r). We illustrate some representative behaviour here, 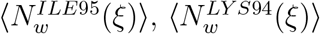 and 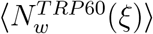 in Fig.6(f). As the ligand goes further from the protein, 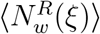 increases around ILE95 and LYS94. Fitting the data with ∼ *ξ*^*α*^ dependence as shown in Fig.6(f), we find that *α*=0.69 for ILE95 and *α*=0.21 for LYS94. The difference in *α* value is due to the fact that LYS94, being basic residue, can interact with water more easily. In contrast, TRP60, which acts as one of the binding residues of the protein but not as an essential coordinate, does not show any change. Our data suggest that the hydrophobic residue, like ILE95, release more water than the other residues, while the ligand approaches the active site of the protein. The larger amount of water release is likely to be aided by the fluctuations in the residue.

The domination of fluctuations can be further observed in the changes of the conformation free energy Δ*G* of ECs in BAMLET with respect to MG-aLA. The free energy change for residue having most EC in BAMLET, namely, ILE 95 is −0.094 kJ/mol for dihedral *ϕ* and −0.97 kJ/mol for dihedral *ψ*. The stabilization of ILE95 is primarily achieved via *ψ*95. Fig.7(a)-(b) shows histogram of free energy change for *ϕ* and *ψ* degree of freedoms (*H*(Δ*G*_*ϕ*_) and *H*(Δ*G*_*ψ*_)) of all the residues in aLA. We find that the residues having ECs, Δ*G* values are clustered near the zero value. Similarly, for *ψ* degree of freedom (Fig.7(b)), most of the residues fall neighborhood zero value. We observe that the residues having EC are clustered near the low change in the thermodynamic parameters, suggesting slow fluctuations. This helps the carries activities in BAMLET state in both ways. The release of water helps efficient binding between the hydrophobic moieties. However, the remnant fluctuations aid the release of the cytotoxic factor. It is known that the inherent dynamic nature of IDPs play a key role in rapid ligand recognition.^40–42^ An IDP protein takes up a stable conformation upon ligand binding.^41,43,44^ However, the fluctuations do not die off in MG state, unlike the case of an IDP.

**Figure 7:**
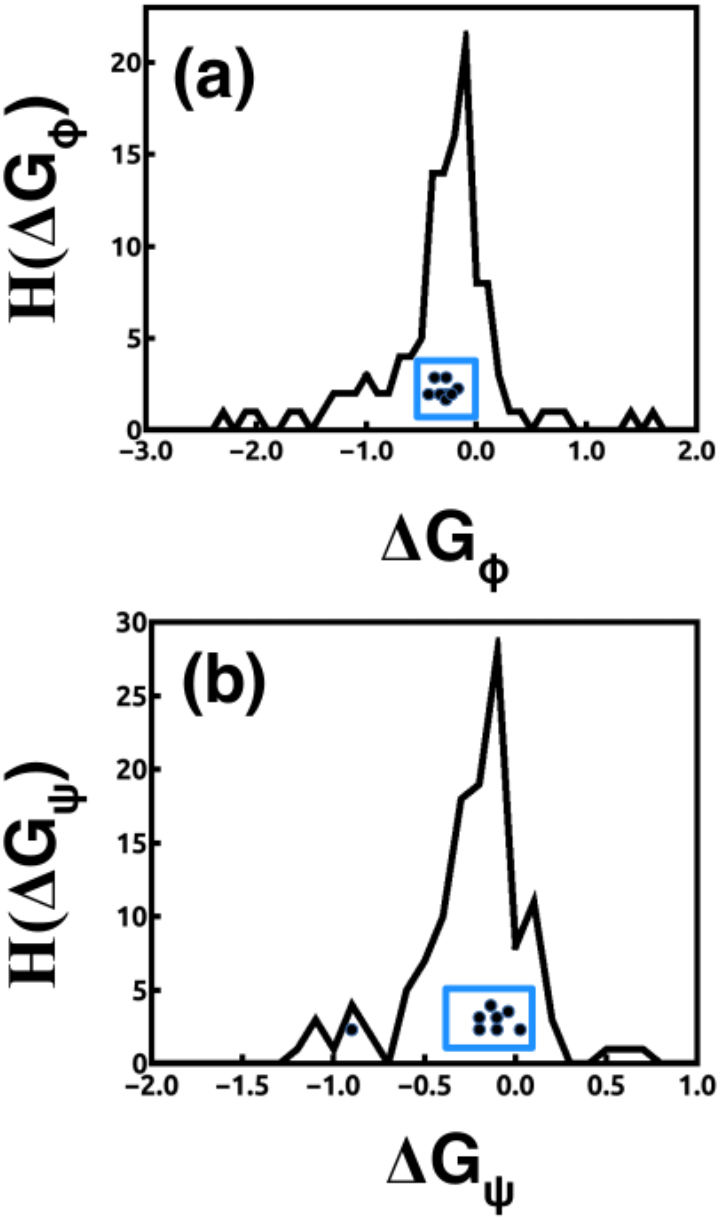
Histogram of Δ*G* value for all residues considering (e) *ϕ* dihedral, (f) *ψ* dihedral angle. Value of Δ*G* for essential residues are marked inside the blue box.

## Conclusion

In conclusion, we show from our analysis on a model for BAMLET that the carrier activity of MG-aLA, involving binding to OLA, is fluctuation dominated. In particular, the residues responsible for binding are also essential for long-lived fluctuations in the system and have low costs of conformation thermodynamic free energy. We expect this scenario to hold for other scenario of ligand binding with protein. A microscopic understanding of fluctuation-driven ligand binding would be helpful in designing drug carrier for targeted drug delivery.

## Supporting information

Suppplementary File

## Acknowledgement

AGM thanks DST India, for an INSPIRE fellowship (IF170961) to conduct the research and Mr. Anirban Paul for helpful discussions.

## Supporting Information Available

**Supporting Table:** Table S1: Docking study on MG-aLA-OLA system. Table S2: Predicted pKa values of titrable residues during CpHMD simulations at pH=2. The offset value is defined as the difference between predicted pKa and system pH. Fraction of time titrable residues remain protonated during simulations. Table S3: Putative binding sites of Oleic acid (OLA) with nature and secondary element in crystal structure.

## References

(1) Ptitsyn, O. B. Protein folding: hypotheses and experiments. Journal of Protein Chemistry 1987, 6, 273–293.

(2) Kuwajima, K. The molten globule state as a clue for understanding the folding and cooperativity of globular-protein structure. Proteins: Structure, Function, and Bioinformatics 1989, 6, 87–103.

(3) Dobson, C. M.; Evans, P. A.; Radford, S. E. Understanding how proteins fold: the lysozyme story so far. Trends in biochemical sciences 1994, 19, 31–37.

(4) Dobson, C. M. Unfolded proteins, compact states and molten globules: Current Opinion in Structural Biology 1992, 2: 6–12. Current Opinion in Structural Biology 1992, 2, 6–12.

(5) Arai, M.; Kuwajima, K. Role of the molten globule state in protein folding. Advances in protein chemistry 2000, 53, 209–282.

(6) Hill, R. L.; Brew, K. Lactose synthetase. Adv Enzymol Relat Areas Mol Biol 1975, 43, 411–490.

(7) Barrick, D.; Baldwin, R. L. The molten globule intermediate of apomyoglobin and the process of protein folding. Protein Science 1993, 2, 869–876.

(8) Jennings, P. A.; Wright, P. E. Formation of a molten globule intermediate early in the kinetic folding pathway of apomyoglobin. Science 1993, 262, 892–896.

(9) Balbach, J.; Forge, V.; van Nuland, N. A.; Winder, S. L.; Hore, P. J.; Dobson, C. M. Following protein folding in real time using NMR spectroscopy. Nature structural biology 1995, 2, 865–870.

(10) Jøhnke, M.; Petersen, T. E. The alpha-lactalbumin/oleic acid complex and its cytotoxic activity. Milk protein 2012,

(11) Barbana, C.; Pérez, M.; Sánchez, L.; Dalgalarrondo, M.; Chobert, J.-M.; Haertlé, T.; Calvo, M. Interaction of bovine α-lactalbumin with fatty acids as determined by partition equilibrium and fluorescence spectroscopy. International dairy journal 2006, 16, 18–25.

(12) Chrysina, E. D.; Brew, K.; Acharya, K. R. Crystal structures of apo-and holo-bovine α-lactalbumin at 2.2-Å resolution reveal an effect of calcium on inter-lobe interactions. Journal of Biological Chemistry 2000, 275, 37021–37029.

(13) Griko, Y. V.; Remeta, D. P. Energetics of solvent and ligand-induced conformational changes in α-lactalbumin. Protein Science 1999, 8, 554–561.

(14) Moulick, A. G.; Chakrabarti, J. Conformational fluctuations in the molten globule state of α-lactalbumin. Phys. Chem. Chem. Phys. 2022, 24, 21348–21357.

(15) Tolin, S.; De Franceschi, G.; Spolaore, B.; Frare, E.; Canton, M.; Polverino de Laureto, P.; Fontana, A. The oleic acid complexes of proteolytic fragments of α-lactalbumin display apoptotic activity. The FEBS journal 2010, 277, 163–173.

(16) Svanborg, C.; Ågerstam, H.; Aronson, A.; Bjerkvig, R.; Düringer, C.; Fischer, W.; Gustafsson, L.; Hallgren, O.; Leijonhuvud, I.; Linse, S., et al. HAMLET kills tumor cells by an apoptosis-like mechanism—cellular, molecular, and therapeutic aspects. Advances in cancer research 2003, 88, 1–29.

(17) Mongan, J.; Case, D. A.; McCammon, J. A. Constant pH molecular dynamics in generalized Born implicit solvent. Journal of computational chemistry 2004, 25, 2038–2048.

(18) Das, A.; Chakrabarti, J.; Ghosh, M. Conformational contribution to thermodynamics of binding in protein-peptide complexes through microscopic simulation. Biophysical journal 2013, 104, 1274–1284.

(19) Sikdar, S.; Chakrabarti, J.; Ghosh, M. A microscopic insight from conformational thermodynamics to functional ligand binding in proteins. Molecular Biosystems 2014, 10, 3280–3289.

(20) Frederick, K. K.; Marlow, M. S.; Valentine, K. G.; Wand, A. J. Conformational entropy in molecular recognition by proteins. Nature 2007, 448, 325–329.

(21) Brandt, S.; Sittel, F.; Ernst, M.; Stock, G. Machine learning of biomolecular reaction coordinates. The journal of physical chemistry letters 2018, 9, 2144–2150.

(22) Sittel, F.; Filk, T.; Stock, G. Principal component analysis on a torus: Theory and application to protein dynamics. The Journal of chemical physics 2017, 147, 244101.

(23) Sittel, F.; Stock, G. Robust density-based clustering to identify metastable conformational states of proteins. Journal of chemical theory and computation 2016, 12, 2426–2435.

(24) Sittel, F.; Stock, G. Perspective: Identification of collective variables and metastable states of protein dynamics. The Journal of chemical physics 2018, 149, 150901.

(25) Swails, J. M.; York, D. M.; Roitberg, A. E. Constant pH replica exchange molecular dynamics in explicit solvent using discrete protonation states: implementation, testing, and validation. Journal of chemical theory and computation 2014, 10, 1341–1352.

(26) Case, D. A.; Cheatham III, T. E.; Darden, T.; Gohlke, H.; Luo, R.; Merz Jr, K. M.; Onufriev, A.; Simmerling, C.; Wang, B.; Woods, R. J. The Amber biomolecular simulation programs. Journal of computational chemistry 2005, 26, 1668–1688.

(27) Hartigan, J. A.; Wong, M. A. Algorithm AS 136: A k-means clustering algorithm. Journal of the royal statistical society. series c (applied statistics) 1979, 28, 100–108.

(28) Roe, D. R.; Cheatham III, T. E. PTRAJ and CPPTRAJ: software for processing and analysis of molecular dynamics trajectory data. Journal of chemical theory and computation 2013, 9, 3084–3095.

(29) Van Zundert, G.; Rodrigues, J.; Trellet, M.; Schmitz, C.; Kastritis, P.; Karaca, E.; Melquiond, A.; van Dijk, M.; De Vries, S.; Bonvin, A. The HADDOCK2. 2 web server: user-friendly integrative modeling of biomolecular complexes. Journal of molecular biology 2016, 428, 720–725.

(30) Honorato, R. V.; Koukos, P. I.; Jiménez-García, B.; Tsaregorodtsev, A.; Verlato, M.; Giachetti, A.; Rosato, A.; Bonvin, A. M. Structural biology in the clouds: the WeNMR-EOSC ecosystem. Frontiers in Molecular Biosciences 2021, 8, 729513.

(31) Van Der Spoel, D.; Lindahl, E.; Hess, B.; Groenhof, G.; Mark, A. E.; Berendsen, H. J. GROMACS: fast, flexible, and free. Journal of computational chemistry 2005, 26, 1701–1718.

(32) Hornak, V.; Abel, R.; Okur, A.; Strockbine, B.; Roitberg, A.; Simmerling, C. Comparison of multiple Amber force fields and development of improved protein backbone parameters. Proteins: Structure, Function, and Bioinformatics 2006, 65, 712–725.

(33) Jorgensen, W. L.; Chandrasekhar, J.; Madura, J. D.; Impey, R. W.; Klein, M. L. Comparison of simple potential functions for simulating liquid water. The Journal of chemical physics 1983, 79, 926–935.

(34) Sprenger, K.; Jaeger, V. W.; Pfaendtner, J. The general AMBER force field (GAFF) can accurately predict thermodynamic and transport properties of many ionic liquids. The Journal of Physical Chemistry B 2015, 119, 5882–5895.

(35) Sousa da Silva, A. W.; Vranken, W. F. ACPYPE-Antechamber python parser interface. BMC research notes 2012, 5, 1–8.

(36) Hub, J. S.; De Groot, B. L.; van der Spoel, D. g wham A Free Weighted Histogram Analysis Implementation Including Robust Error and Autocorrelation Estimates. Journal of chemical theory and computation 2010, 6, 3713–3720.

(37) Swaminathan, S.; Harte Jr, W.; Beveridge, D. L. Investigation of domain structure in proteins via molecular dynamics simulation: application to HIV-1 protease dimer. Journal of the American Chemical Society 1991, 113, 2717–2721.

(38) Chakraborty, K.; Bandyopadhyay, S. Correlated Conformational Motions of the KH Domains of Far Upstream Element Binding Protein Complexed with Single-Stranded DNA Oligomers. The Journal of Physical Chemistry B 2015, 119, 10998–11009.

(39) Grant, B. J.; Rodrigues, A. P.; ElSawy, K. M.; McCammon, J. A.; Caves, L. S. Bio3d: an R package for the comparative analysis of protein structures. Bioinformatics 2006, 22, 2695–2696.

(40) Uversky, V. N. Intrinsic disorder, protein–protein interactions, and disease. Advances in protein chemistry and structural biology 2018, 110, 85–121.

(41) Wright, P. E.; Dyson, H. J. Linking folding and binding. Current opinion in structural biology 2009, 19, 31–38.

(42) Wright, P. E.; Dyson, H. J. Intrinsically disordered proteins in cellular signalling and regulation. Nature reviews Molecular cell biology 2015, 16, 18–29.

(43) Sugase, K.; Dyson, H. J.; Wright, P. E. Mechanism of coupled folding and binding of an intrinsically disordered protein. Nature 2007, 447, 1021–1025.

(44) Van Der Lee, R.; Buljan, M.; Lang, B.; Weatheritt, R. J.; Daughdrill, G. W.; Dunker, A. K.; Fuxreiter, M.; Gough, J.; Gsponer, J.; Jones, D. T., et al. Classification of intrinsically disordered regions and proteins. Chemical reviews 2014, 114, 6589–6631.

