## Supplementary material for "Fluctuation dominated ligand binding in molten globule protein": Suppplementary File

### Supplementary Information article "Fluctuation dominated ligand binding in molten globule protein"

Table S1: Docking study on BAMLET system.

| | |
|---|---|
| HADDOCK score | -34.4 +/- 2.4 |
| Cluster size | 19 |
| RMSD from the overall lowest-energy structure | 1.9 +/- 0.0 |
| Van der Waals energy | -21.8 +/- 1.1 |
| Electrostatic energy | -93.6 +/- 1.4 |
| Desolvation energy | -6.7 +/- 0.2 |
| Restraints violation energy | 34.9 +/- 19.2 |
| Buried Surface Area | 613.0 +/- 11.9 |
| Z-Score | -2.0 |

Table S2: Predicted pKa values of titrable residues during CpHMD simulations at pH=2. The offset value is defined as the difference between predicted pKa and system pH. Fraction of time titrable residues remain protonated during simulations.

| Residue Name | Residue | Predicted pKa Value | Offset Value | Fraction of protonation |
|---|---|---|---|---|
| 7 | GLU | 3.675 | 1.675 | 0.956 |
| 11 | GLU | 3.334 | 1.334 | 0.987 |
| 14 | GLU | 3.869 | 1.869 | 0.794 |
| 25 | GLU | 2.586 | 0.586 | 0.998 |
| 37 | ASP | 3.239 | 1.239 | 0.945 |
| 46 | ASP | 3.652 | 1.652 | 0.978 |
| 49 | GLU | 3.933 | 1.933 | 0.988 |
| 63 | ASP | 2.472 | 0.472 | 0.748 |
| 64 | ASP | 3.270 | 1.270 | 0.949 |
| 78 | ASP | 2.714 | 0.714 | 0.838 |
| 82 | ASP | 2.594 | 0.594 | 0.797 |
| 83 | ASP | 3.077 | 1.077 | 0.923 |
| 84 | ASP | 2.784 | 0.784 | 0.859 |
| 87 | ASP | 2.052 | 0.052 | 0.530 |
| 88 | ASP | 3.499 | 1.499 | 0.969 |
| 97 | ASP | 3.392 | 1.392 | 0.961 |
| 113 | GLU | 4.661 | 2.661 | 0.998 |
| 116 | ASP | 3.282 | 1.282 | 0.950 |
| 121 | GLU | 4.091 | 2.091 | 0.992 |

Table S3: Putative binding sites of Oleic acid (OLA) with nature and secondary element in crystal structure.

| Residue Number | Residue | Initial Structure | Nature |
|---|---|---|---|
| 51 | GLY | Loop | Hydrophobic |
| 52 | LEU | Loop | Hydrophobic |
| 53 | PHE | Loop | Hydrophobic |
| 55 | ILE | Sheet | Hydrophobic |
| 59 | ILE | Loop | Hydrophobic |
| 60 | TRP | Loop | Hydrophobic |
| 89 | ILE | Helix | Hydrophobic |
| 90 | MET | Helix | Hydrophobic |
| 92 | VAL | Helix | Hydrophobic |
| 93 | LYS | Helix | Basic |
| 94 | LYS | Helix | Basic |
| 95 | ILE | Helix | Hydrophobic |
| 96 | LEU | Helix | Hydrophobic |
| 98 | LYS | Helix | Basic |
| 99 | VAL | Helix | Hydrophobic |
| 100 | GLY | Helix | Hydrophobic |
| 101 | ILE | Helix | Hydrophobic |
| 104 | TRP | Helix | Hydrophobic |
